# SEX-SPECIFIC TRAJECTORIES OF REELIN-DEPENDENT MATURATION OF DEEP LAYER PREFRONTAL NEURONS

**DOI:** 10.1101/2021.12.23.473977

**Authors:** Thenzing Judá Silva Hurtado, Olivier Lassalle, Antoine Ameloot, Pascale Chavis

## Abstract

Throughout early adulthood, the anatomical and functional maturation of PFC circuitry continues under the influence of multiple extrinsic and intrinsic factors, most notably electrical activity, and molecular cues. We previously showed that the extracellular matrix protein reelin orchestrates the structural and functional maturation of deep layers medial PFC (mPFC) pyramidal neurons. Additionally, we reported that reelin haploinsufficiency is associated to prefrontal disruptions of long-term memory retention thereby illustrating the eminent role of reelin in cognitive maturation of the PFC. Prefrontal maturation follows a sex-specific developmental pattern, supporting the existence of sexual differences in the morphology-functional properties PFC. Here, we interrogated the role of reelin in the functional maturation of excitatory networks in the mPFC. The developmental trajectory of reelin’s expression and deep layer pyramidal neurons synaptic plasticity was tracked in the mPFC of male and female mice, from the juvenile period to adulthood. To assess the role of reelin in both sexes, wild-type and heterozygous reeler mice (HRM) were compared. The results show that the maturational profile of reelin expression in the mPFC is sex-dependent and that the developmental trajectory of long-term potentiation is different between wild-type males and females. These data demonstrate reelin’s influence on prefrontal synapses is sex and period specific.

## INTRODUCTION

Reelin is a signaling glycoprotein serving multiple functions in the brain throughout life which has also emerged as a psychiatric risk factor in a wide spectrum of psychiatric disorders (Jossin 2020). Secreted by Cajal-Retzius cells in the marginal zone of the cerebral cortex and hippocampus during embryonic development, reelin plays an essential role in neuronal migration, positioning and layer formation (Sekine et al. 2014). In addition to being a developmental molecule, reelin is an important contributor to postnatal and adult central nervous system (CNS) physiology. In postnatal forebrain, reelin production is shifted to subpopulations of GABAergic interneurons distributed throughout cellular layers of the hippocampus and neocortex (Alcantara et al. 1998; Pesold et al. 1998; Campo et al. 2009). In the maturing and adult CNS, reelin modulates several aspects of excitatory and inhibitory synaptic function and morpho-functional plasticity. Reelin plays an important role in dendritic maturation and spine development (Niu et al. 2008; Chameau et al. 2009), in the maturation of NMDA receptors (Sinagra et al. 2005; Groc et al. 2007; Campo et al. 2009), hippocampal long-term potentiation, synaptic transmission and cognitive ability (Weeber et al. 2002; Beffert et al. 2005; Pujadas et al. 2010; Rogers et al. 2011). Data from our laboratory have shown that reelin is necessary for the correct structural and functional maturation of excitatory synapses in the prelimbic area of the prefrontal cortex (PL-PFC). Spine density, excitatory synaptic transmission and plasticity of deep prefrontal pyramidal neurons as well as cognitive traits are altered during the postnatal maturation of reelin-haploinsufficient heterozygous reeler mice PL-PFC (Iafrati et al. 2014; Iafrati et al. 2016). We have also shown that reelin is necessary for the fine-tuning of GABAergic synaptic transmission in deep layer prefrontal pyramidal neurons as well as for the physiological excitatory/inhibitory balance in the maturing PL-PFC (Bouamrane et al. 2017). Most of these studies were performed on male progeny. Despite these advancements, it remains unknown whether the role of reelin in the physiology of the postnatal CNS is identical in both sexes.

The PFC is an associative brain region that supports complex cognitive functions. One distinctive feature of the PFC is a protracted maturation into adulthood (Gogtay et al. 2004) characterized by intense synaptogenesis, remodeling of dendritic spines and connectivity refinement in parallel to maturation of cognitive abilities (van Eden et al. 1990; Petanjek et al. 2011). The protracted PFC maturation is also under the influence of hormonal factors and several maturational processes, such as fear discrimination (Day et al. 2020), frontal white matter growth (Willing and Juraska 2015), parvalbumin neurons density (Du et al., 2018) and pyramidal cells complexity (Markham et al. 2013), were reported to follow a sex-specific developmental pattern. Collectively, these evidence show that pubertal onset and sex are important determinants of PFC postnatal maturation.

In the present study, we undertook a cross-sectional study to explore the maturational trajectory of reelin expression in the PL-PFC and of excitatory synaptic plasticity in layer V PL-PFC pyramidal neurons from male and female mice. We used electrophysiological and immunohistochemical analyses from the juvenile period to adulthood in wild-type mice and heterozygous reeler mice (HRM). We discovered a sexual divergence in the developmental trajectory of reelin expression, associated to an abrupt increase in the density of reelin-positive neurons in adult wild-type males in contrast to a stable reelin expression across maturation in wild-type females. Our data also uncovered a delayed expression of long-term potentiation (LTP) in wild-type females and marked sex-specific differences in the effect of reelin haploinsufficiency on LTP.

## MATERIALS AND METHODS

### Animals

Heterozygous reeler mice (HRM) and their wild-type littermate were bred by intercrossing males and females HRM from the B6C3Fe a/a-Relnrl/J strain (Jackson Laboratory). Offsprings were genotyped by PCR as previously described (Iafrati et al., 2014). All mice were weaned at 21 days and then caged socially in same-sex groups. Mice were housed in standard 12 h light–dark cycle and supplied food pellets and water *ad libitum*. Animals were treated in strict compliance with the criteria of the European Communities Council Directive (agreement number 2015121715284829-V4).

### Electrophysiology

Coronal slices containing the prelimbic area (PL) of the medial prefrontal cortex (mPFC) were prepared as previously described (Iafrati et al., 2014). Briefly, mice were anesthetized with isoflurane and 300 μm-thick coronal slices were prepared in a sucrose-based solution at 4°C using an Integraslice vibratome (Campden Instruments). Slices were stored for 30 min at 32°C in artificial cerebrospinal fluid (ACSF) containing (in mM): NaCl (130), KCl (2.5), MgCl_2_ (2.4), CaCl_2_ (1.2), NaHCO_3_ (23), NaH_2_PO_4_ (1.2) and Glucose (11), equilibrated with 95% O2 /5% CO2. Slices were then stored at room temperature until recording. All experiments were conducted at 30-32°C in ACSF. For recording, slices were superfused at 2 ml per min with ACSF containing Picrotoxin (100µM; Sigma) or SR95531 (Gabazine, 5µM; Tocris 1262) to block GABA_A_ receptors.

The prelimbic area of the mPFC was visualized using an infrared illuminated upright microscope (Olympus BX51WI, France) and extracellular field excitatory postsynaptic potentials (fEPSPs) recordings carried out as previously described (Iafrati et al., 2014; 2016). fEPSPs were recorded in the PL-PFC layer V with an ACSF-filled electrode and evoked in layer III at 0.1 Hz with a stimulating glass electrode filled with ACSF. LTP was induced using a theta-burst stimulation protocol consisting of five trains of burst with four pulses at 100 Hz, at 200 ms interval, repeated four times at intervals of 10 s. The glutamatergic nature of fEPSPs was confirmed at the end of each experiment by perfusing the non-NMDA (N-methyl-D-aspartate) ionotropic glutamate receptor antagonist 6-cyano-7-nitroquinoxaline-2,3-dione (CNQX) (20 μM; NIH), which specifically blocked the synaptic component without altering the non-synaptic component (not shown)

Signals were collected using an Axopatch-200B amplifier (Axon Instruments, Molecular Devices, Sunnyvale, USA), filtered at 2 kHz, digitized at 10 kHz, acquired with Clampex 10.7 acquisition Software via a Digidata 1440A (Axon Instruments) and analyzed using Axograph 1.7.6.

### Immunohistochemistry

Animals were deeply anesthetized with Pentobarbital (90mg/kg; Exagon Med’Vet) and perfused transcardially with cold phosphate-buffered saline solution (PBS; Gibco Life Technologies) followed with Antigenfix (DiaPath) a phosphate-buffered paraformaldehyde solution. The dissected brains were postfixed overnight at 4°C in the same fixative. Brains were then sectioned using a vibratome (VT 1200 s, Leica) into 60 μm-thick coronal slices.

Sections were first rinsed three times for 10 min in PBS and then incubated in blocking solution with 0.1M PBS containing 0.3% Triton X100 (Sigma) and 0.2% Bovine Serum Albumin (BSA; Sigma) twice for 1h at room temperature (RT). Slices were incubated free-floating overnight at RT with the mouse G10 anti-reelin primary antibody (1:3000, MAB5406; Millipore) diluted in the blocking solution. After three blocking solution washes (10 min each), section was incubated at RT for 75 min with the secondary antibody donkey anti-mouse Alexa 568 (Invitrogen ThermoFisher Scientific) diluted 1:500 in the blocking solution. After three rinses of 10 min in 0.1 M PBS, sections were stained with Hoechst (Invitrogen ThermoFisher Scientific) diluted 1:1000 in 0.1M PBS for 12 min, washed again three times for 10 min in 0.1 M PBS and coverslipped with Aqua-Poly/Mount (Polysciences).

The specificity of the G10 antibody was tested on sections obtained from homozygous reeler mice lacking reelin expression. Additional negative controls were performed by omitting the G10 primary antibody on wild-type or HRM sections.

### Image analysis

Confocal images were acquired with a Zeiss LSM-800 system equipped with emission spectral detection and a tunable laser providing excitation range from 470 to 670 nm. Stacks of optical sections were collected with a Plan Apochromat 20x.0.8 air objective for 3D reconstructions. Laser power and photomultiplier gain were adjusted to obtain few pixels with maximum intensity on the somata showing the higher labelling intensity. To obtain a whole rostro-caudal representation of the PL-PFC of each mouse, images were acquired from Bregma 2.58 to 1.94 according to the Mouse Brain Atlas (Paxinos & Franklin 2009). To obtain a Z-representation across layers I to VI within each brain section, scanning was performed using tiles representing a total of 894.4 μm x 894.4 μm surface/image size and a Z-stack selection covering a depth of 17.5 to 19.8 µm. The tri-dimensional reconstruction and blind semiautomated analysis were performed with Fiji (Image J).

### Statistical Analysis

All values are given as mean ± SEM and statistical significance was set at *P* < 0.05. Statistical analysis was performed with GraphPad Prism 9.2.0 (GraphPad Software, La Jolla, CA, USA). Two sample comparisons were made with the non-parametric Mann–Whitney test and multiple comparisons were made using a one-way analysis of variance (ANOVA) followed, if significant, by Tukey’s multiple comparisons test.

## RESULTS

Reelin expression and excitatory synaptic transmission were analyzed in PL-PFC throughout the first 4 months of postnatal life following the maturational epochs delineated in our previous reports: juvenile period (Juv, P22-28), adolescence (Ado, P30-45) and adulthood (Adu, P70-115; Iafrati et al. 2016; Bouamrane et al. 2017). We focused on layer V pyramidal neurons, one of the main output cells of the PL-PFC microcircuit.

### Maturational trajectory of reelin expressing neurons in the PL-PFC of wild-type male and female mice

The expression of reelin was traced in the PL-PFC of wild-type males and females from the juvenile period until adulthood (Figure 1). In males, the density of reelin-expressing neurons in the whole PL-PFC was stable in juvenile and adolescents and increased abruptly during adulthood, whereas in females it remained stable across maturation (Figure 1A). We also examined whether the sex-dependent maturational profile of reelin expression was mirrored in the different layers of the PL-PFC. We found that reelin expression followed the same pattern in layers I, II/III and V/VI, with an increased density in adult males (Figure 1B) and a stable expression at all maturational stages in females (Figure 1D).

**Figure 1:**
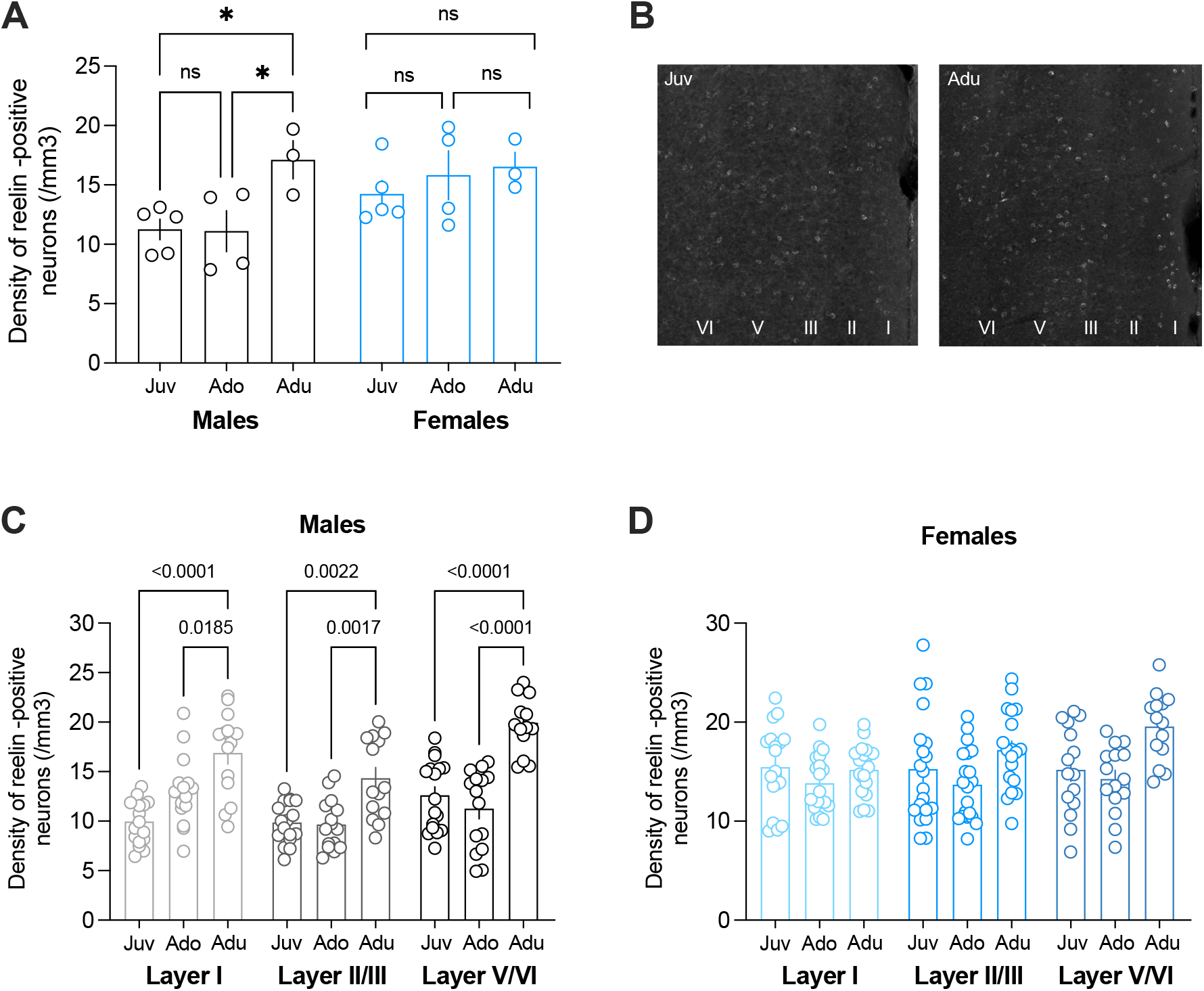
Sex-dependent developmental profile of reelin expression in the PL PFC of wild-type mice. **A:** Number of reelin-positive neurons per mm3 in the PL-PFC of wild-type male and female mice across development. In males, the average (± sem) density of reelin-positive neurons was 11.2 ± 0.9 in juvenile mice (n=5; Juv, P22-26) and 11.1 ± 1.7 in adolescent mice (n=4; Ado, P35-40) and increase to 17.1 ± 1.6 in adults (n=3; Adu, P95-100; F_(2,9)_=5.397, P=0.0288, ANOVA). In females, the average density of reelin-positive neurons remained stable from juvenile (14.2 ± 1.1, n=5; Juv, P22-26), to adolescent (15.8 ± 2.0, n=4; Ado, P35-40) and adults (16.5 ± 1.2, n=3; Adu, P115; F_(2,9)_=0.597, P=0.5709, ANOVA). n represents the number of animals. **B:** Confocal images of PL-PFC sections stained for reelin from Juvenile (P25) and Adult (P94) male wild-type mice. **C:** Developmental trajectory of reelin-expressing neurons across PLPFC layers I, II/III and V/VI in wild-type male mice. In all layers, the density of reelin-positive neurons was stable until adolescence and increased sharply at adulthood (P values indicated on the bar graph, ANOVA). **D:** In wild-type female mice, the density of reelin-expressing neuron was similar within layers I, II/II and V/VI across maturation. C-D: n indicates the number of slices selected in a rostro-caudal axis for each mouse.

These data show that the maturational trajectory of reelin expression is sex but not layer dependent in wild-type mice. In the context of our previous findings showing that reelin levels influence the morpho-functional maturation of deep layer PL-PFC pyramidal neurons, we next examined whether the maturational trajectory of long-term potentiation (LTP) and effects of reelin haploinsufficiency on LTP were sex-dependent.

### Sex-specific developmental sequence of TBS long-term potentiation in wild type mice

In wild-type male mice, theta-burst stimulation (TBS) induced a long-term potentiation (LTP) of field excitatory postsynaptic potentials (fEPSP) measured at synapses onto layer V pyramidal neurons. This TBS-LTP was robustly expressed from the juvenile period to adulthood (Figure 2D and Supplementary Figure 1A). In contrast, in juvenile wild-type females TBS-LTP was markedly reduced compared to juvenile males (Figure 2A and 2D). During adolescence and adulthood, the time course and magnitude of TBS-LTP in females PL-PFC are similar to age-matched male mice (Figure 2B-D). These results show that the expression of full-fledged LTP is delayed by a month in wild-type females (Supplementary Figure 1B).

**Figure 2:**
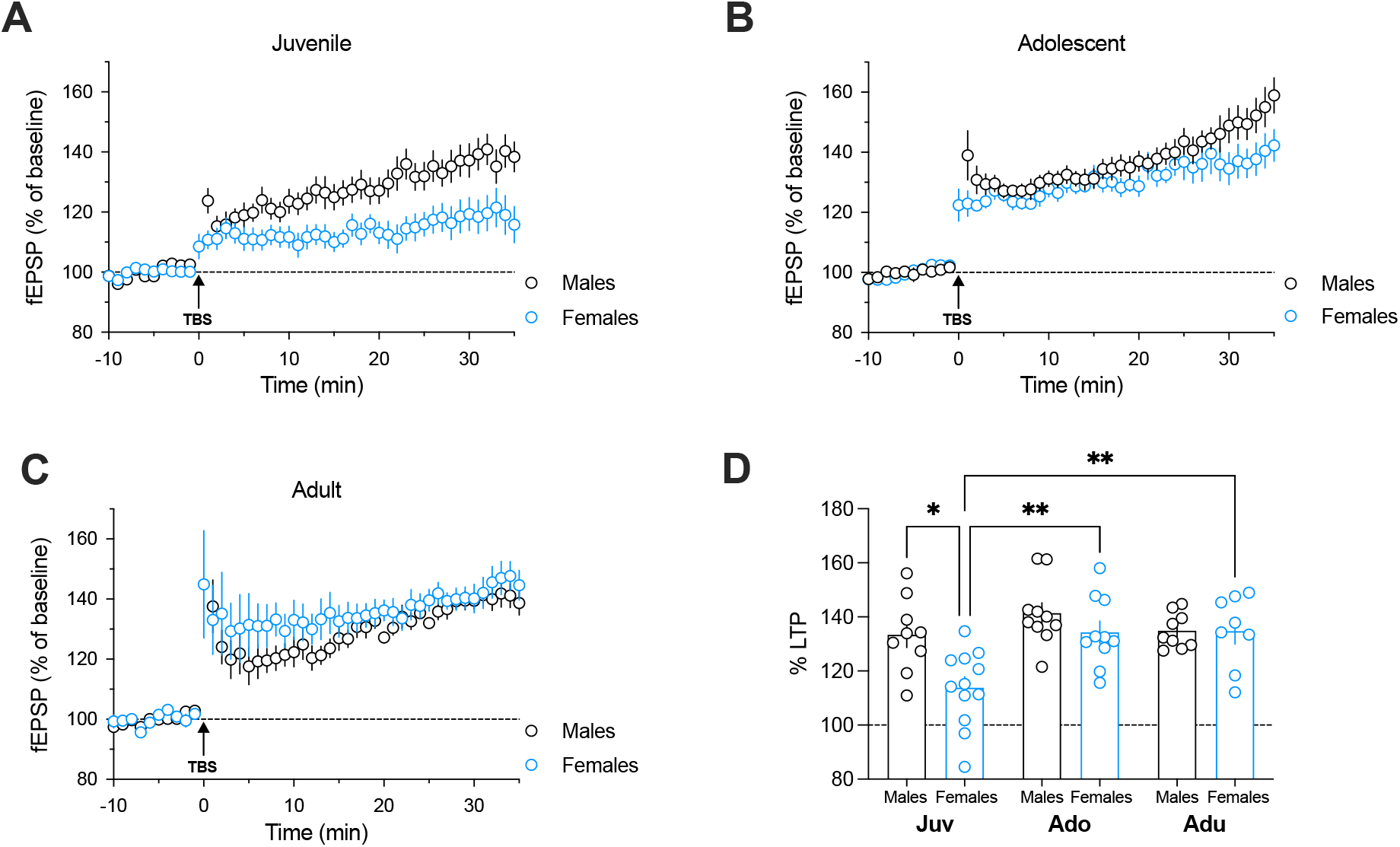
Sex-dependent developmental profile of synaptic plasticity in layer V of PLPFC in wild-type mice. **A:** Grouped time courses of field excitatory postsynaptic potential (fEPSP) responses expressed as the percentage of baseline before and after theta-burst stimulation (TBS, indicated by arrow). During the juvenile period (Juv, P22-28), TBS-induced LTP of PL-PFC synapses in wild-type females (n=12) was reduced compared to male mice (n=9). **B:** Grouped time courses of fEPSP amplitudes (percentage of baseline before and after TBS) showing similar TBS-LTP in both males (n=10) and females (n=10) during adolescence (Ado, P30-45). **C:** In adults (Adu, P70-115), the time course of normalized fEPSP responses (percentage of baseline before and after TBS) are similar in males (n=9) and females (n=8). **D:** The percentage of potentiation was measured 20-30 min after TBS. Juvenile males: 133.4 ± 4.6% (n=9), females: 113.8 ± 4.1% (n=12); Adolescent males: 141.4 ± 3.8% (n=10), females: 134.3 ± 4.1% (n=10); Adult males: 134.9 ± 2.2% (n=9), females: 134.7 ± 4.8% (n=8). F_(5,52)_=6.419, P=0.0001, ANOVA.* P<0.05, **P<0.01. All data are means ± sem.

The data indicate that in males, TBS-LTP is mature at the juvenile stage and consistently expressed until adulthood, whereas in females, LTP fully develops at adolescence. The delay in the maturation profile of LTP between males and female show a sex-dependent maturational trajectory of TBS-LTP in PL-PFC deep layers.

### Sex-specific effect of reelin haploinsufficiency on long-term potentiation

Recently, we have shown that TBS-LTP is altered in the PL-PFC of the reelin-haploinsufficient heterozygous reeler mice (HRM; Iafrati et al., 2014; Iafrati et al., 2016), but whether this effect is different in males and females is not known. To test this hypothesis, we followed the developmental profile of TBS-LTP from juvenile to adults in males and females HRM (Figures 3 & 4).

**Figure 3:**
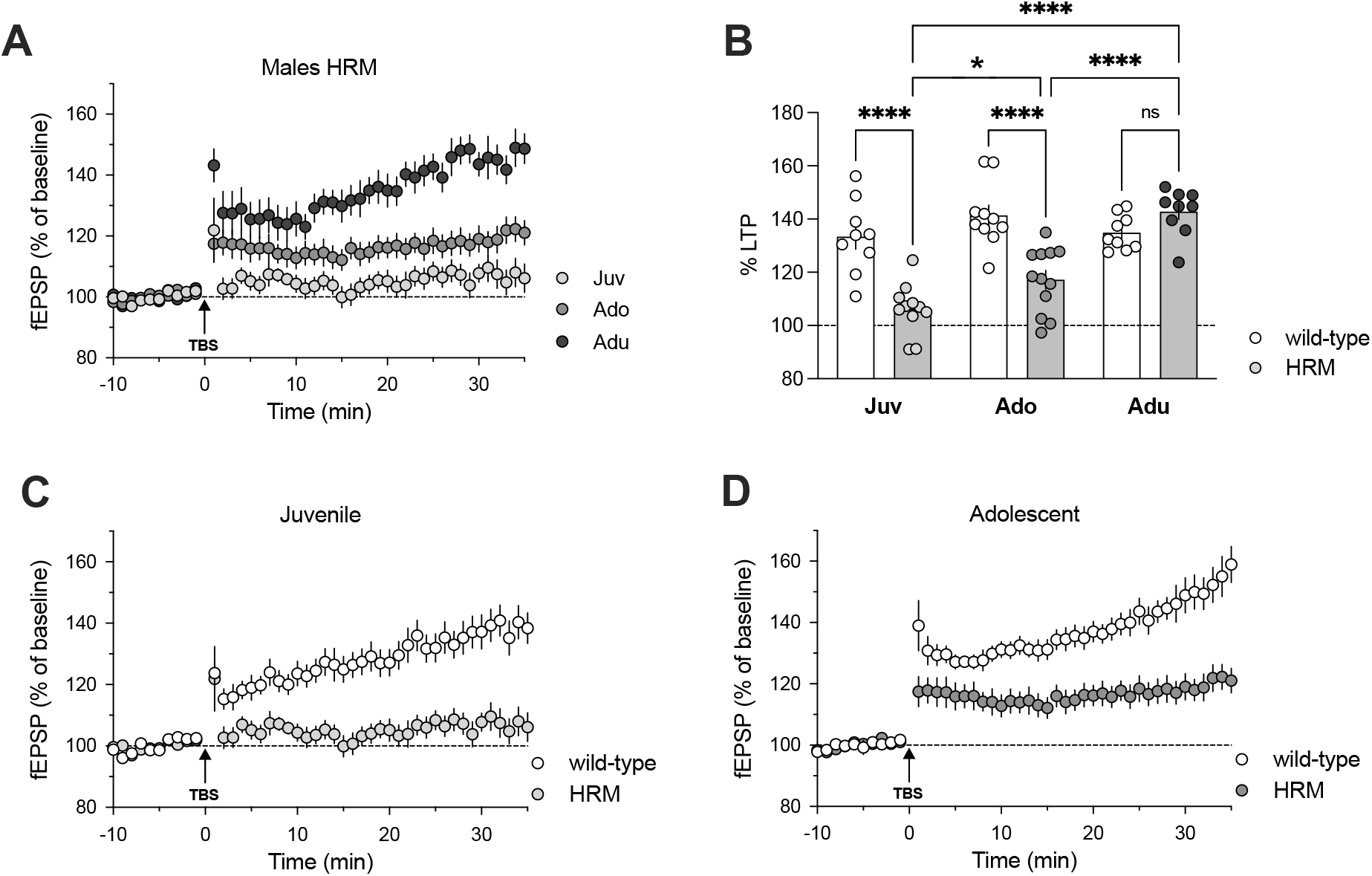
Effect of reelin haploinsufficiency on the developmental profile of TBS-LTP in male mice. **A:** Grouped time courses of fEPSP responses (percentage of baseline before and after TBS) in males HRM showing that TBS-LTP gradually develops from juvenile (n=11) to adult period (n=9). **B:** The percentage of potentiation measured 20-30 min after LTP induction was: 133.4 ± 4.6% (n=9) in wild-type and 106.2 ± 2.8% (n=11) in HRM during the juvenile period, 141.4 ± 3.8% (n=10) and 117.2 ± 3.5% (n=12) in adolescent wild-type and HRM respectively, and during adulthood 134.9 ± 2.2% (n=9) for wild-type and 142.8 ± 2.9% (n=9) for HRM. F_(5,54)_=18.85, P<0.0001, ANOVA. **C-D:** Average time courses of mean fEPSPs amplitudes before and after TBS (percentage of baseline) showing the absence of LTP in male HRM during the juvenile period (**C**) and a reduced LTP during adolescence in HRM compared to wild-type mice (**D**). * P<0.05, ****P<0.0001. All data are means ± sem.

In males HRM, plasticity gradually developed throughout the juvenile and adolescent periods (Figure 3A). TBS-LTP was abolished in juvenile HRM (Figure 3B-C), markedly reduced during adolescence (Figure 3B, 3D) and robustly expressed in adults HRM compared to age-matched wild-type males (Figure 3B & Supplementary Figure 2). In adults, time course and magnitude of TBS-LTP were similar in wild-type mice and HRM (Figure 3B & Supplementary Figure 2). The downregulation of plasticity by reelin haploinsufficiency could not be attributed to differences in the excitability of layer III/V excitatory synapses as input-output curves were superimposable between wild-type and HRM across all ages (Supplementary Figure 3).

In contrast to juvenile males HRM lacking TBS-LTP, TBS induced a LTP in juvenile females HRM that was undistinguishable from LTP obtained in aged-matched wild-type females (Supplementary Figure 4A & Figure 4B). Unlike wild-type and HRM males during adolescence, TBS-LTP was similar in both genotypes in females (Figure 4B & Supplementary Figure 4B). Until adolescence, no differences were observed in the excitability of layer III/V excitatory synapses between females wild-type and HRM (Supplementary Figure 4C-D). In adults HRM females, TBS-LTP was largely reduced compared to age-matched wild-type mice (Figure 4B-C). This effect was correlated to a marked reduction of synaptic excitability in females HRM compared to wild-type mice (Figure 4D). This is in marked contrast to what was observed in males at the same age (Supplementary Figure 2).

**Figure 4:**
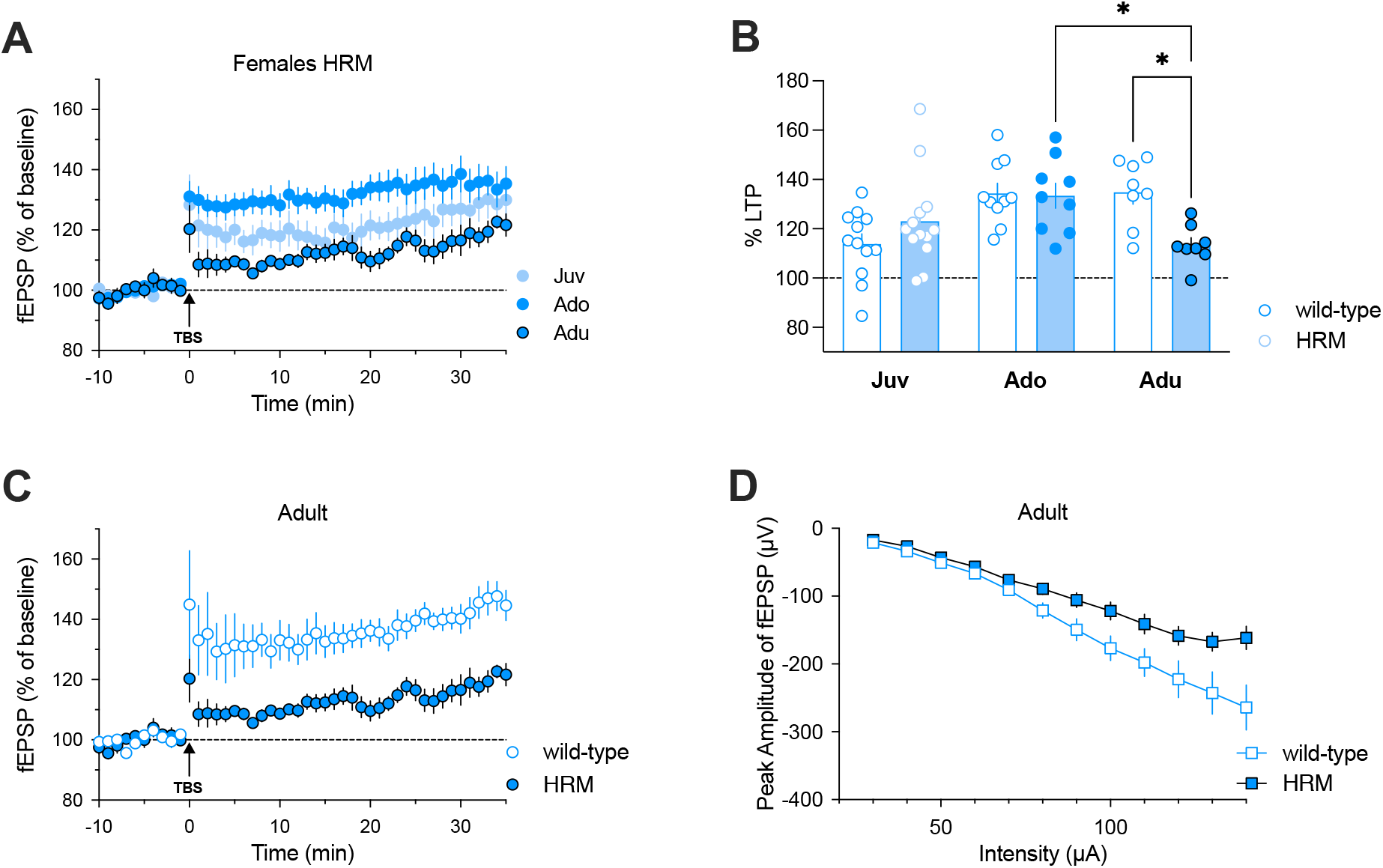
Effect of reelin haploinsufficiency on the developmental profile of TBS-LTP in female mice. **A:** Time course of normalized fEPSP responses (percentage of baseline before and after TBS) in female HRM at each developmental period showing that in adults TBS-LTP decreases to levels similar to those measured in juvenile HRM (Juv: n=13; Ado: n=9; Adu: n=8). **B:** Bar graph of fEPSP percentage change from baseline 20-30 min after TBS. Juvenile wild-type: 113.8 ± 4.1% (n=12) and HRM 123.0 ± 5.2% (n=13); adolescent wild-type: 134.3 ± 4.1% (n=10) and HRM: 133.4 ± 5.0 (n=9); adult wild-type: 134.7 ± 4.8% (n=8) and HRM: 113.5 ± 2.9% (n=8). F_(5,54)_=4.569, P=0.0015, ANOVA. **C:** Average time courses of normalized fEPSPs amplitudes before and after TBS showing a reduced TBS-LTP in HRM (n=8) compared to wild-type mice (n=8) at the adult stage. **D:** Input-ouput curves of average fEPSP amplitudes in response to stimulation intensity showing that the excitability of layer III/V synapses is reduced in adult HRM (n=8) compared to wild-type mice (n=8). *P<0.05. All data are means ± sem.

These data show that the effects of decreased levels of reelin on TBS-LTP occur at different maturational epochs in males and females. All together, these results show that the disruption of TBS-LTP at PL-PFC layers III/V synapses by reelin haploinsufficiency is age- and sex-specific.

## DISCUSSION

This cross-sectional study shows that in deep layer (layer V) pyramidal neurons of the mouse PL-PFC, the developmental profile of reelin expression, the maturation of excitatory input TBS-induced synaptic plasticity, and the effect of reelin haploinsufficiency on LTP follow a sex-specific trajectory from the juvenile period until adulthood. Specifically, we showed that while TBS-LTP is already matured in juvenile males, in females it fully develops during adolescence. We also showed an age-related and sex-specific disruption of TBS-LTP by reelin haploinsufficiency. Whereas in males HRM LTP impairment was observed from the juvenile period through adolescence, in female HRM alteration of LTP was observed only during adulthood.

The mechanisms underlying these effects are sex-dependent as synaptic excitability measured by the input-output relationship was reduced only in adult females HRM but not in juvenile or adolescent males HRM. The reduction of the LTP in the HRM could also result from an alteration of various mechanisms known to be modulated by treelin, such as the probability of glutamate release (Hellwig et al. 2011), the morpho-functional maturation of the excitatory synapses, the repertoire of the ionotropic receptors of the AMPA and NMDA type glutamate (Iafrati et al. 2016), the signaling pathways involved in LTP. We previously reported in HRM mice that the magnitude of the LTP was strongly correlated with the AMPA / NMDA ratio and the dendritic spines (Iafrati et al. 2016). Whether the alteration of LTP in females HRM could be attributable to a morpho-functional alteration of the excitatory synapses generated by inadequate reelin levels during the maturation of glutamatergic circuits is not known.

Although estradiol can be metabolized in males from cholesterol, this sex hormone is predominantly produced cyclically by ovaries. Estradiol is involved in a multitude of mechanisms that can impact the LTP. Indeed, this hormone improves synaptic transmission by facilitating the insertion of AMPA receptors to synaptic sites and increases the LTP in the hippocampus (Kramar et al. 2013). In addition, estradiol increases BDNF levels, a growth factor also promoting LTP (Hill & van den Buuse 2011). During postnatal maturation, female mice present more microglia than males (Hanamsagar & Bilbo 2016). Microglia could play a role in the mechanisms of synaptic plasticity mechanisms during development, in particular by interacting with the extracellular matrix and participating in remodeling and elimination of dendritic spines following synaptic activity (Hanamsagar & Bilbo 2016). In HRM male mice, the alteration of the LTP has been associated with morphological defects of dendritic spines (Iafrati et al. 2016), making this last mechanism an interesting candidate to explain the sexual differences of the pathological developmental trajectories of the PL-PFC.

A significant contribution of reelin to the etiology of psychiatric and neurodevelopmental disorders has been proposed based on evidence of the roles of reelin in adult and developing brain together with patients’ data showing alteration in reelin levels (Folsom and Fatemi 2013). Patients suffering from psychiatric disorders such as schizophrenia, bipolar disorder, major depression, and autism spectrum disorders (ASD) exhibit an approximate reelin downregulation of 50% in several brain structures, most notably the hippocampus and the PFC (Impagnatiello et al. 1998; Guidotti et al. 2000; Folsom and Fatemi 2013). Several studies suggest that alterations of the postnatal PFC maturation may contribute to the development of psychiatric disorders including depression, addiction, ASD and schizophrenia (Lewis 1997; Raedler et al. 1998).

Noteworthy, in human sex-specific differences also observed in the expression of psychiatric disorders with schizophrenic males having an earlier onset than females. The present data provide a possible mechanism underlying the differential vulnerabilities to psychiatric disorders exiting between sexes.

## Author contributions

TJSH performed experiments and analyzed the data. OL and AA performed electrophysiology and analyzed the data. PC supervised the project, analyzed data and wrote the paper.

## Acknowledgements

The authors are grateful to A. Montheil and F. Bader of the INMED genotyping facility and thank the members of INMED animal facility. This work was supported by the Institut National de la Santé et de la Recherche Médicale (INSERM), Fondation Jerôme Lejeune (R13139AA), CONACYT (339066/471725, TJSH). We thank the National Institute of Mental Health’s Chemical Synthesis and Drug Supply Program (Rockville, MD, USA) for providing CNQX.

---

**Supplementary Figure 1:**
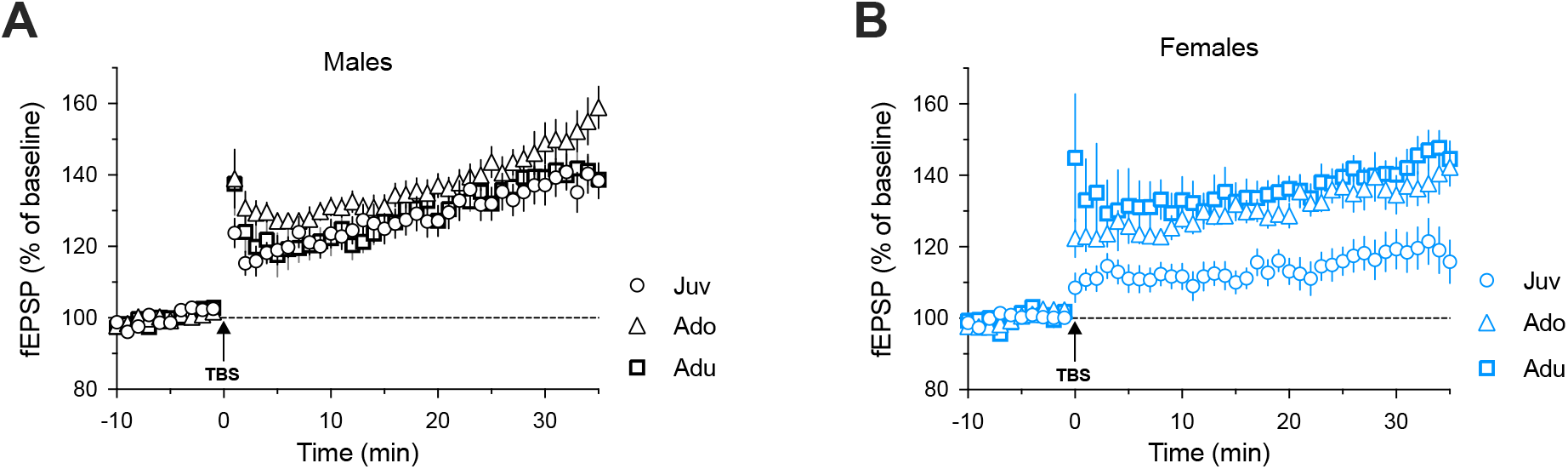
Developmental trajectory of TBS-induced LTP within each sex. **A:** Grouped time courses representing the mean ± sem of layer III/V fEPSPs expressed as the percentage of baseline before and after TBS in wild-type male mice across maturation. TBS-LTP remained similar in all age groups (Juv, n=9 mice; Ado, n=10; Adu, n=9 mice). **B:** Average (± sem) time courses of fEPSP amplitudes (percentage of baseline before and after TBS) in juvenile (n=12), adolescent (n=10) and adult (n=8) female mice. The magnitude of TBS-LTP was lower during the juvenile period compared to adolescence and adulthood.

**Supplementary Figure 2:**
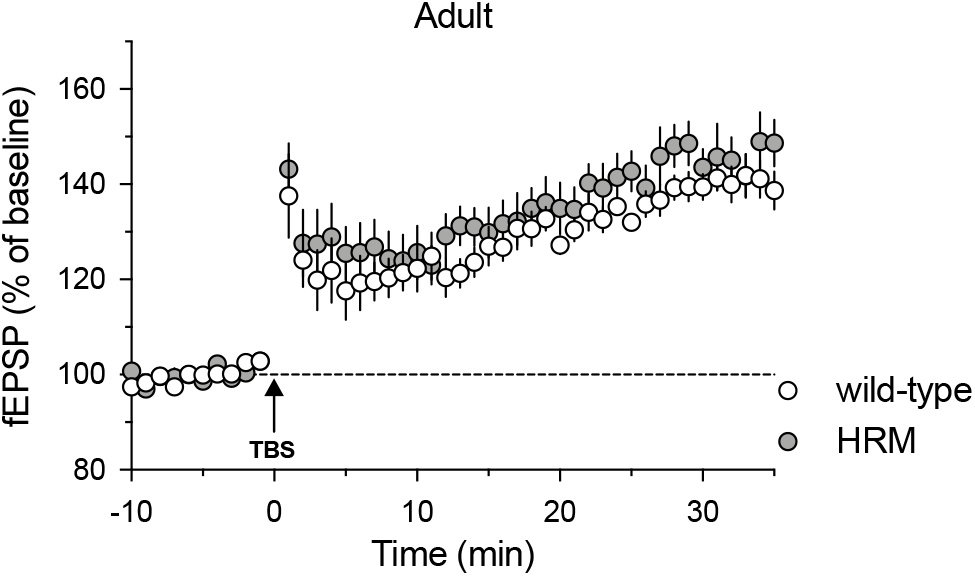
TBS-LTP at layer III/V synapses is robustly expressed in both males wild-type mice and HRM at adulthood. Average time course of mean ± sem fEPSPs amplitudes before and after TBS (percentage of baseline) in both genotypes (wild-type: n=9, HRM: n=9).

**Supplementary Figure 3:**
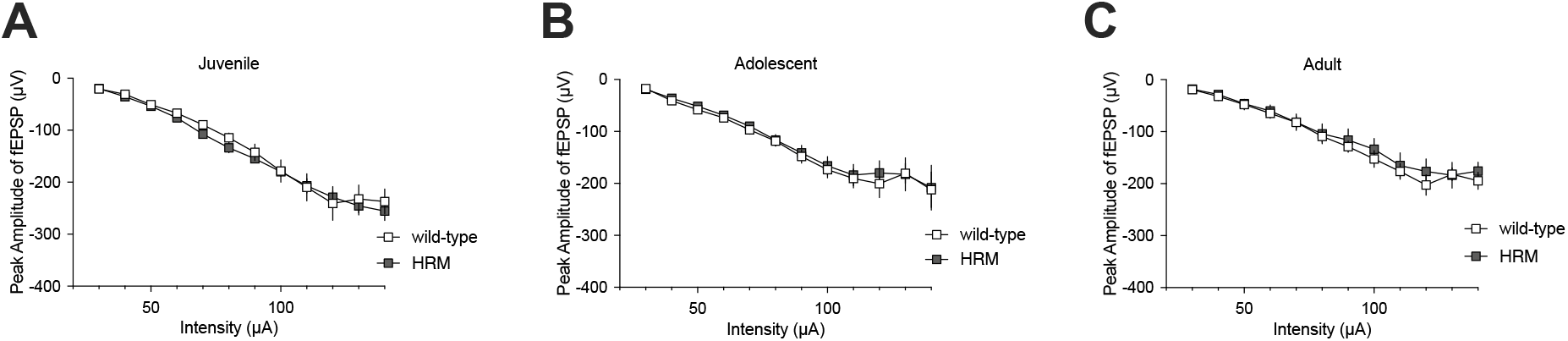
Excitability of layer III/V synapses is not affected by reelin-haploinsufficiency across maturation in male mice. Input-output profile of average fEPSP amplitude (± sem) is shown for wild-type mice versus HRM at each developmental period. Juvenile: n=9 wild-type and n=11 HRM; adolescent n=10 wild-type and n=12 HRM; adult: n=9 wild-type and n=9 HRM.

**Supplementary Figure 4:**
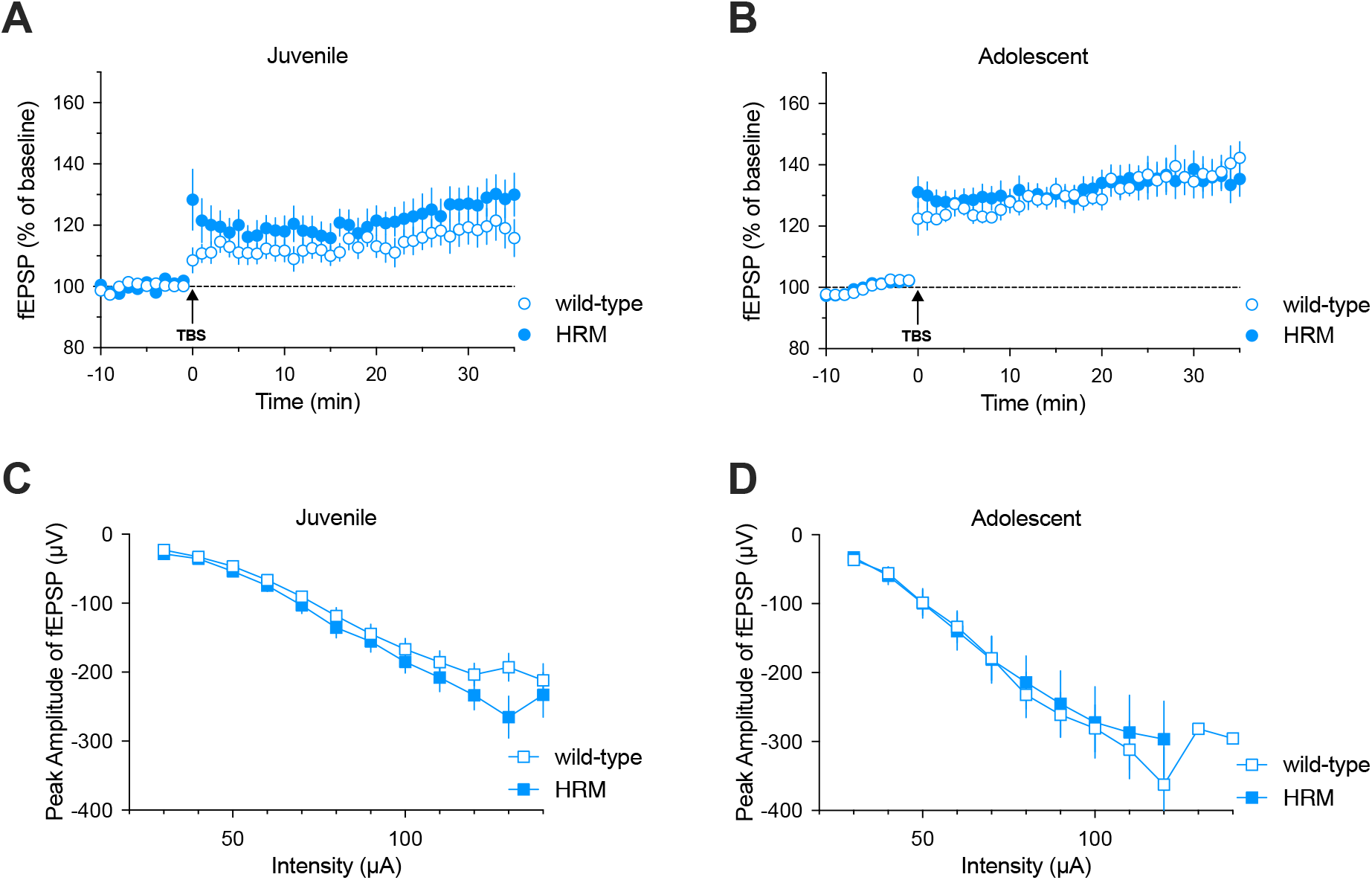
Plasticity and excitability of layer III/V synapses are comparable between wild-type and HRM females in juvenile and adolescent periods. **A-B:** Time course of TBS-LTP normalized to baseline in wild-type mice and HRM during the juvenile period (**A**) and adolescence (**B**). **C-D:** Input-output curves of the average fEPSP amplitudes recorded in layer V in response to stimulation intensity in layer III in juvenile wild-type (n=12) and HRM (n=13; **C**) and adolescent wild-type (n=10 and HRM (n=9; **D**). All data are means ± sem.

## REFERENCES

Alcantara, S., Ruiz, M., D’Arcangelo, G., Ezan, F., de Lecea, L., Curran, T., et al. (1998). Regional and cellular patterns of reelin mRNA expression in the forebrain of the developing and adult mouse. J. Neurosci 18, 7779–7799.

Beffert, U., Weeber, E. J., Durudas, A., Qiu, S., Masiulis, I., Sweatt, J. D., et al. (2005). Modulation of synaptic plasticity and memory by Reelin involves dierential splicing of the lipoprotein receptor Apoer2. Neuron 47, 567–579.

Bouamrane L, Scheyer AF, Lassalle O, Iafrati J, Thomazeau A, Chavis P. Reelin-Haploinsufficiency Disrupts the Developmental Trajectory of the E/I Balance in the Prefrontal Cortex. Front Cell Neurosci. 2017 Jan 12;10:308. doi: 10.3389/fncel.2016.00308. eCollection 2016.

Campo, C. G., Sinagra, M., Verrier, D., Manzoni, O. J., and Chavis, P. (2009). Reelin secreted by GABAergic neurons regulates glutamate receptor homeostasis. PLoS ONE 4:e5505. doi: 10.1371/journal.pone.0005505

Chameau, P., Inta, D., Vitalis, T., Monyer, H., Wadman, W. J., and van Hooft, J. A. (2009). The N-terminal region of reelin regulates postnatal dendritic maturation of cortical pyramidal neurons. Proc. Natl. Acad. Sci. U.S.A. 106, 7227–7232. doi: 10.1073/pnas.0810764106

Day, H.L., Suwansawang, S., Halliday, D.M. & Stevenson, C.W. (2020) Sex differences in auditory fear discrimination are associated with altered medial prefrontal cortex function. Sci. Rep. 10, 6300.

Du X, Serena K, Hwang WJ, Grech AM, Wu YWC, Schroeder A, Hill RA. 2018. Prefrontal cortical parvalbumin and somatostatin expression and cell density increase during adolescence and are modified by BDNF and sex. Molecular and Cellular Neuroscience 88:177–188. DOI: https://doi.org/10.1016/j.mcn.2018.02.001, PMID: 29408239

Folsom, T. D., and Fatemi, S. H. (2013). The involvement of Reelin in neurodevelopmental disorders. Neuropharmacology 68, 122–135. doi: 10.1016/j.neuropharm.2012.08.015

Gogtay, N., Giedd, J. N., Lusk, L., Hayashi, K. M., Greenstein, D., Vaituzis, A. C., et al. (2004). Dynamic mapping of human cortical development during childhood through early adulthood. Proc. Natl. Acad. Sci. U.S.A. 101, 8174– 8179. doi: 10.1073/pnas.0402680101

Groc, L., Choquet, D., Stephenson, F. A., Verrier, D., Manzoni, O. J., and Chavis, P. (2007). NMDA receptor surface tracking and synaptic subunit composition are developmentally regulated by the extracellular matrix protein Reelin. J. Neurosci. 27, 10165–10175. doi: 10.1523/JNEUROSCI.1772-07.2007

Guidotti, A., Auta, J., Davis, J. M., Di-Giorgi-Gerevini, V., Dwivedi, Y., Grayson, D. R., et al. (2000). Decrease in reelin and glutamic acid decarboxylase67 (GAD67) expression in schizophrenia and bipolar disorder: a postmortem brain study. Arch. Gen. Psychiatry 57, 1061–1069. doi: 10.1001/archpsyc.57.11.1061

Hanamsagar, R., & Bilbo, S. D. (2016). Sex differences in neurodevelopmental and neurodegenerative disorders: focus on microglial function and neuroinflammation during development. The Journal of steroid biochemistry and molecular biology, 160, 127–133.

Hellwig, S. et al., 2011. Role for Reelin in Neurotransmitter Release. Journal of Neuroscience, 31(7), pp.2352–2360.

Hill, R. A., & van den Buuse, M. (2011). Sex-dependent and region-specific changes in TrkB signaling in BDNF heterozygous mice. Brain research, 1384, 51–60.

Iafrati, J., Malvache, A., Gonzalez Campo, C., Orejarena, M. C., Lassalle, O., Bouamrane, O., et al. (2016). Multivariate synaptic and behavioral profiling reveals new developmental endophenotypes in the prefrontal cortex. Sci. Rep. 6, 35504. doi: 10.1038/srep35504

Iafrati, J., Orejarena, M. J., Lassalle, O., Bouamrane, L., Gonzalez-Campo, C., and Chavis, P. (2014). Reelin, an extracellular matrix protein linked to early onset psychiatric diseases, drives postnatal development of the prefrontal cortex via GluN2B-NMDARs and the mTOR pathway. Mol. Psychiatry 19, 417–426. doi: 10.1038/mp.2013.66

Impagnatiello, F., Guidotti, A. R., Pesold, C., Dwivedi, Y., Caruncho, H., Pisu, M. G., et al. (1998). A decrease of reelin expression as a putative vulnerability factor in schizophrenia. Proc. Natl. Acad. Sci. U.S.A. 95, 15718–15723. doi: 10.1073/pnas.95.26.15718

Kramár, E. A., et al. (2013). Estrogen promotes learning-related plasticity by modifying the synaptic cytoskeleton. Neuroscience, 239, 3–16.

Lewis, D. A. (1997). Development of the prefrontal cortex during adolescence: insights into vulnerable neural circuits in schizophrenia. Neuropsychopharmacology 16, 385–398. doi: 10.1016/S0893-133X(96)00277-1

Markham J.A., Mullins S.E., Koenig J.I. Periadolescent maturation of the prefrontal cortex is sex-specific and is disrupted by prenatal stress. J Comp Neurol. 2013 Jun 1;521(8):1828–43. doi: 10.1002/cne.23262.

Niu, S., Yabut, O., and D’Arcangelo, G. (2008). The Reelin signaling pathway promotes dendritic spine development in hippocampal neurons. J. Neurosci. 28, 10339–10348. doi: 10.1523/JNEUROSCI.1917-08.2008

Pesold, C., Impagnatiello, F., Pisu, M. G., Uzunov, D. P., Costa, E., Guidotti, A., et al. (1998). Reelin is preferentially expressed in neurons synthesizing gamma-aminobutyric acid in cortex and hippocampus of adult rats. Proc. Natl. Acad. Sci. U.S.A. 95, 3221–3226. doi: 10.1073/pnas.95.6.3221

Petanjek, Z. et al. Extraordinary neoteny of synaptic spines in the human prefrontal cortex. Proceedings of the National Academy of Sciences of the United States of America 108, 13281– 13286 (2011).

Pujadas, L., Gruart, A., Bosch, C., Delgado, L., Teixeira, C. M., Rossi, D., et al. (2010). Reelin regulates postnatal neurogenesis and enhances spine hypertrophy and long-term potentiation. J. Neurosci. 30, 4636–4649. doi: 10.1523/JNEUROSCI.5284-09.2010

Raedler, T. J., Knable, M. B., and Weinberger, D. R. (1998). Schizophrenia as a developmental disorder of the cerebral cortex. Curr. Opin. Neurobiol. 8, 157–161. doi: 10.1016/S0959-4388(98)80019-6

Sekine, K., Kubo, K., and Nakajima, K. (2014). How does Reelin control neuronal migration and layer formation in the developing mammalian neocortex? Neurosci. Res. 86, 50–58. doi: 10.1016/j.neures.2014.06.004

Sinagra, M., Verrier, D., Frankova, D., Korwek, K. M., Blahos, J., Weeber, E. J., et al. (2005). Reelin, very-low-density lipoprotein receptor, and apolipoprotein E receptor 2 control somatic NMDA receptor composition during hippocampal maturation in vitro. J. Neurosci. 25, 6127– 6136. doi: 10.1523/JNEUROSCI.1757-05.2005

van Eden, C. G., Kros, J. M. & Uylings, H. B. The development of the rat prefrontal cortex. Its size and development of connections with thalamus, spinal cord and other cortical areas. Progress in brain research 85, 169–183 (1990).

Weeber, E. J., Beert, U., Jones, C., Christian, J. M., Forster, E., Sweatt, J. D., et al. (2002). Reelin and ApoE receptors cooperate to enhance hippocampal synaptic plasticity and learning. J. Biol. Chem. 277, 39944–39952. doi: 10.1074/jbc.M205147200

Willing, J and Juraska J.M. The timing of neuronal loss across adolescence in the medial prefrontal cortex of male and female rats. Neuroscience 301 (2015) 268–275

